# Chemical inhibition of pathological reactive astrocytes promotes neural protection

**DOI:** 10.1101/2021.11.03.467083

**Authors:** Benjamin L.L. Clayton, James D. Kristell, Kevin C. Allan, Molly Karl, Eric Garrison, Yuka Maeno-Hikichi, Annalise M. Sturno, H. Elizabeth Shick, Robert H. Miller, Paul J. Tesar

**Affiliations:** Department of Genetics and Genome Sciences, Case Western Reserve University School of Medicine, Cleveland, OH, USA; Department of Anatomy and Regenerative Biology, George Washington University School of Medicine, Washington D.C., USA

## Abstract

Disease, injury, and aging induce reactive astrocyte states with pathological functions^1-4^. In neurodegenerative diseases, inflammatory reactive astrocytes are abundant and contribute to progressive cell loss. Modulating the state or function of these reactive astrocytes thereby represents an attractive therapeutic goal^5,6^. Leveraging a cellular phenotypic screening platform, we show that chemical inhibitors of HDAC3 effectively block pathological astrocyte reactivity. Inhibition of HDAC3 reduces molecular and functional features of reactive astrocytes *in vitro* including inflammatory gene expression, cytokine secretion, and antigen presentation. Transcriptional and chromatin mapping studies show that HDAC3 inhibition mediates a switch between pro-inflammatory and anti-inflammatory states, which disarms the pathological functions of reactive astrocytes. Systemic administration of a blood-brain barrier penetrant chemical inhibitor of HDAC3, RGFP966, blocks reactive astrocyte formation and promotes axonal protection *in vivo*. Collectively, these results establish a platform for discovering chemical modulators of reactive astrocyte states, inform the mechanisms controlling astrocyte reactivity, and demonstrate the therapeutic potential of modulating astrocyte reactivity for neurodegenerative diseases.

## Main

Astrocytes in the central nervous system (CNS) play important homeostatic roles that include trophic support of neurons, promotion of functional synapse formation, and formation and maintenance of the blood-brain barrier^7-9^. In the context of disease, injury, or normal aging, astrocytes become reactive and can adopt a pathological state that kills neurons and oligodendrocytes^6^. These pathogenic reactive astrocytes are found in neurodegenerative diseases including Alzheimer’s disease, Parkinson’s disease, Huntington’s disease, amyotrophic lateral sclerosis, vanishing white matter disease^10^, and multiple sclerosis^1,11^. Thus, there is considerable interest in modulating pathological reactive astrocytes to reduce the progression of these diseases^3,6,12^. Here, we develop an astrocyte discovery platform and leverage the power of high-throughput phenotypic drug screening^13^ to identify modulators of reactive astrocytes. We identify HDAC3 as a druggable target that modulates reactive astrocytes *in vitro* and *in vivo* by suppressing pro-inflammatory gene programs while inducing expression of anti-inflammatory genes and genes associated with a beneficial reactive astrocyte state. Using a CNS-penetrant HDAC3 inhibitor, we further show that HDAC3 inhibition decreases the formation of reactive astrocytes and that suppression of pathological reactive astrocytes through HDAC3 inhibition promotes axonal protection *in vivo*.

### An astrocyte discovery platform for high-throughput phenotypic drug screening

Current astrocyte isolation protocols are limited by scale or confounded by culture conditions containing serum, which irreversibly alters the astrocyte transcriptome and morphology^14^. This challenges their use as cellular platforms for high-throughput drug screening. To overcome this, we developed an approach to isolate and culture large numbers (hundreds of millions) of resting mouse cortical astrocytes that exhibit a prototypical stellate morphology and express canonical astrocyte marker proteins including GFAP, AQP4, GLT-1, and ALDH1L1 (Extended Data Fig. 1a,b). Using single-cell RNA sequencing and *in situ* hybridization, we demonstrated the high purity of these cultures and confirmed that the astrocytes become reactive in response to microglial-derived cytokines tumor necrosis factor (TNF), interleukin 1 alpha (IL1α), and complement component 1q (C1q)^1^ (Fig. 1a-c). Comparison to publicly available single-cell RNA sequencing data from systemic lipopolysaccharide (LPS) driven neuroinflammation in mice demonstrated that these reactive astrocytes increase expression of genes that define an *in vivo* neuroinflammatory specific astrocyte state (Extended Data Fig. 1c), which is also found in mouse models of Alzheimer’s disease, multiple sclerosis, and stab wound injury^3^. Additional comparison to astrocyte transcriptional signatures from single-nuclei analysis of Alzheimer’s disease^15^, Huntington’s disease^16^, and Parkinson’s disease^17^ patient tissue showed that our reactive astrocytes reflect the pathological reactive astrocyte states found in these human neurodegenerative diseases (Fig. 1d-f). Finally, we confirmed that our reactive astrocyte cultures acquire functions of reactive astrocytes including cross-presentation of antigen on major histocompatibility complex (MHC) Class I and increased secretion of proinflammatory cytokines^18^ (Fig. 1g,h). Together, these results show that this culture system generates pure astrocytes that robustly transition to a disease-relevant state.

**Figure 1.**
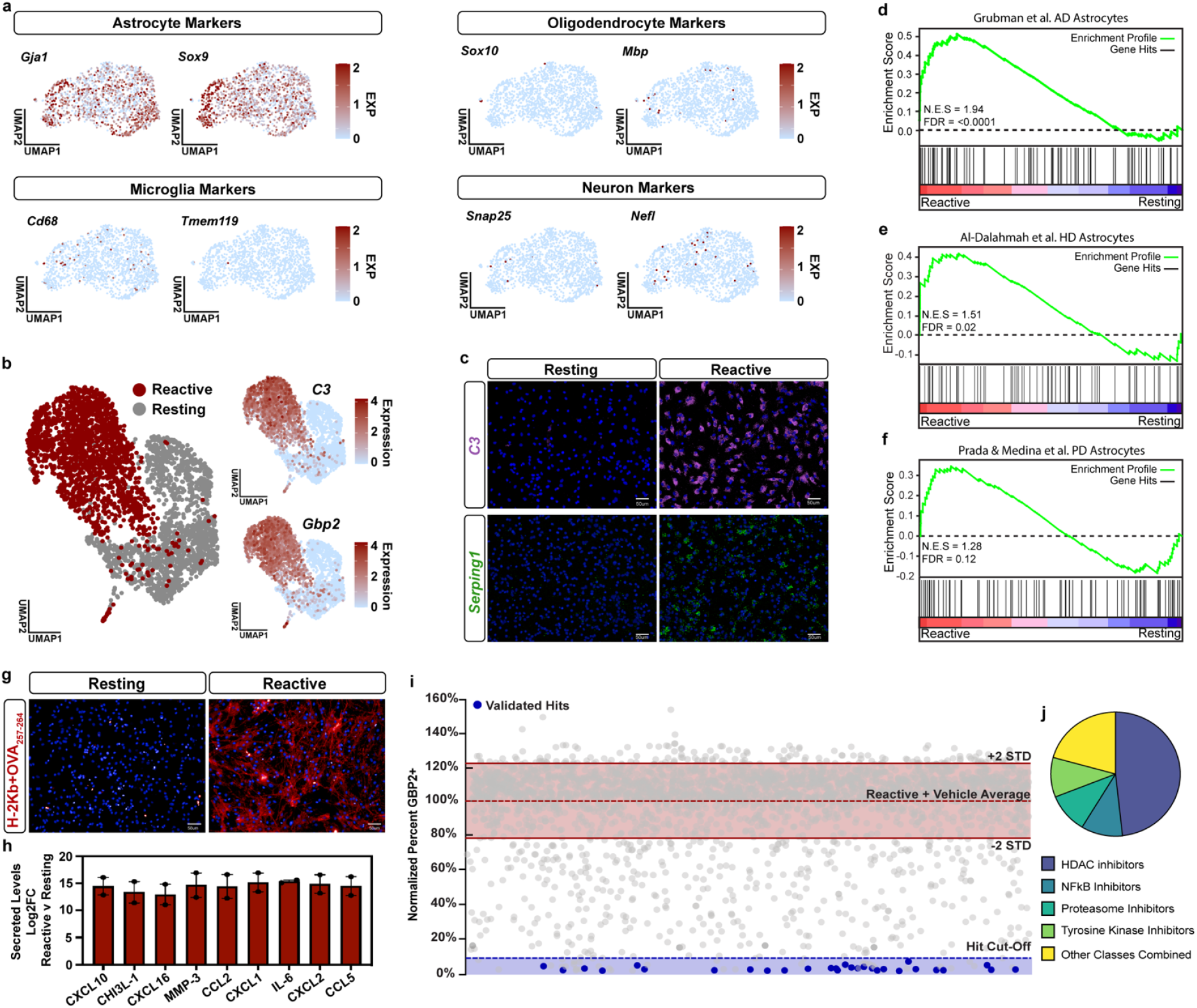
An astrocyte discovery platform identifies inhibitors of pathological reactive astrocyte formation. **a**, Expression of cell-type specific markers layered onto UMAP plots from single-cell RNAseq of primary resting astrocytes. **b**, UMAP plot of resting and pathological reactive astrocytes (abbreviated to reactive in figures) single-cell RNAseq colored by condition, resting (grey) and reactive (red). Additional UMAP plots showing expression levels for the pathological reactive astrocyte markers *C3* and *Gbp2*. **c**, Representative images of *in situ* hybridization in resting and reactive astrocyte cultures with probes against the pathological reactive astrocyte markers *C3* and *Serping1*. **d-f**, Gene set enrichment analysis (GSEA) comparing pathological reactive astrocytes to the top 100 genes upregulated in astrocytes from single-nuclei RNAseq data from **d**, Alzheimer’s, **e**, Huntington’s, and **f**, Parkinson’s disease patient tissue. **g**, Representative images of resting and reactive astrocyte cultures exposed to the OVA_257-264_ peptide and then stained for MHC Class I bound to OVA_257-264_ (H-2Kb+OVA_257-264_) in red. **h**, The Log2 fold-change (Log2FC) of secreted cytokines in reactive vs resting astrocytes conditioned media. Data presented as mean ± s.e.m for *n* = 2 biological replicates (independent astrocyte isolations). **i**, Scatter plot of primary screen results displayed as percent GBP2 positive, normalized to reactive astrocyte plus vehicle controls for all non-toxic chemicals and validated hit chemicals colored in blue. The dashed blue line represents the hit cut-off at a ≥90% decrease in GBP2-positive astrocytes compared to reactive astrocyte plus vehicle controls. Dashed red line represents the average percent GBP2 positive for reactive astrocytes plus vehicle set at 100%. Solid lines represent +/- 2 standard deviations from the mean of reactive plus vehicle control wells. **j**, Pie chart depicting the chemical class breakdown of all 29 validated chemical hits.

### Phenotypic drug screen identifies small-molecules that suppress pathological reactive astrocytes

Leveraging our astrocyte discovery platform we used the power of high-throughput phenotypic screening to identify small-molecules that suppress astrocyte reactivity. Resting astrocytes exposed to TNF, IL1α, and C1q were treated with 3115 small molecules in a 384-well plate format and, after 24 hours, their transition to reactive astrocytes was measured with the marker protein guanylate-binding protein 2 (GBP2), which is specifically upregulated in both mouse and human reactive astrocytes in a pathological state^1,19^. The assay displayed consistently high signal-to-background ratio across all screening plates, demonstrating its suitability as a phenotypic endpoint (Extended Data Fig. 2a-c). Our primary screen successfully identified hit compounds that decreased the percentage of GBP2-positive astrocytes by ≥90% compared to cultures treated with DMSO vehicle (Fig. 1i). Of those primary hits, 29 small molecules were validated to modulate reactive astrocytes over at least 3 concentrations and decrease the expression of proteasome 20S subunit beta 8 (*Psmb8*) as a secondary marker of pathological reactive astrocytes (Extended Data Fig. 2d,e). Validated hits covered a broad range of compound classes but were highly enriched for histone deacetylase (HDAC) inhibitors that accounted for 48% (14/29) of the hits (Fig. 1j and 2a). There are 11 HDAC isozymes and many chemical HDAC inhibitors are non-specific. Only HDAC3 was a shared target of all 14 HDAC inhibitors identified in our primary screen, including the only isozyme-specific chemical, RGFP966 (Extended Data Fig. 3a). RGFP966 is an HDAC3 specific inhibitor with no reported inhibition of other HDAC isozymes at up to 15uM^20^. This led us to hypothesize that HDAC3 was the HDAC inhibitor target responsible for modulating pathological reactive astrocytes.

### HDAC3 regulates pathological reactive astrocyte gene expression and function

Using an additional HDAC3 specific inhibitor, T247^21^, and genetic ablation of *Hdac3*^*22,23*^, we pharmacologically and genetically confirmed that HDAC3 inhibition modulates pathological reactive astrocytes (Extended Data Fig. 3b-f). Inhibition of HDAC3 in human astrocyte cultures also suppressed astrocyte reactivity showing that the effects of HDAC3 inhibition are conserved across species (Extended Data Fig. 3g). RNA sequencing also confirmed that RGFP966 treatment significantly downregulated expression of multiple defining genes of pathological reactive astrocytes, including *C3* and *Serping1*, showing that the effects were global and not limited to decreasing *Gbp2* (Fig. 2b). Pharmacological inhibition of HDAC3 by RGFP966 also suppressed the MHC Class I antigen cross-presentation and inflammatory cytokine secretion functions of pathological reactive astrocytes (Fig. 2c-e). These data show that HDAC3 inhibition reduces pathological reactive astrocyte gene expression and function.

**Figure 2.**
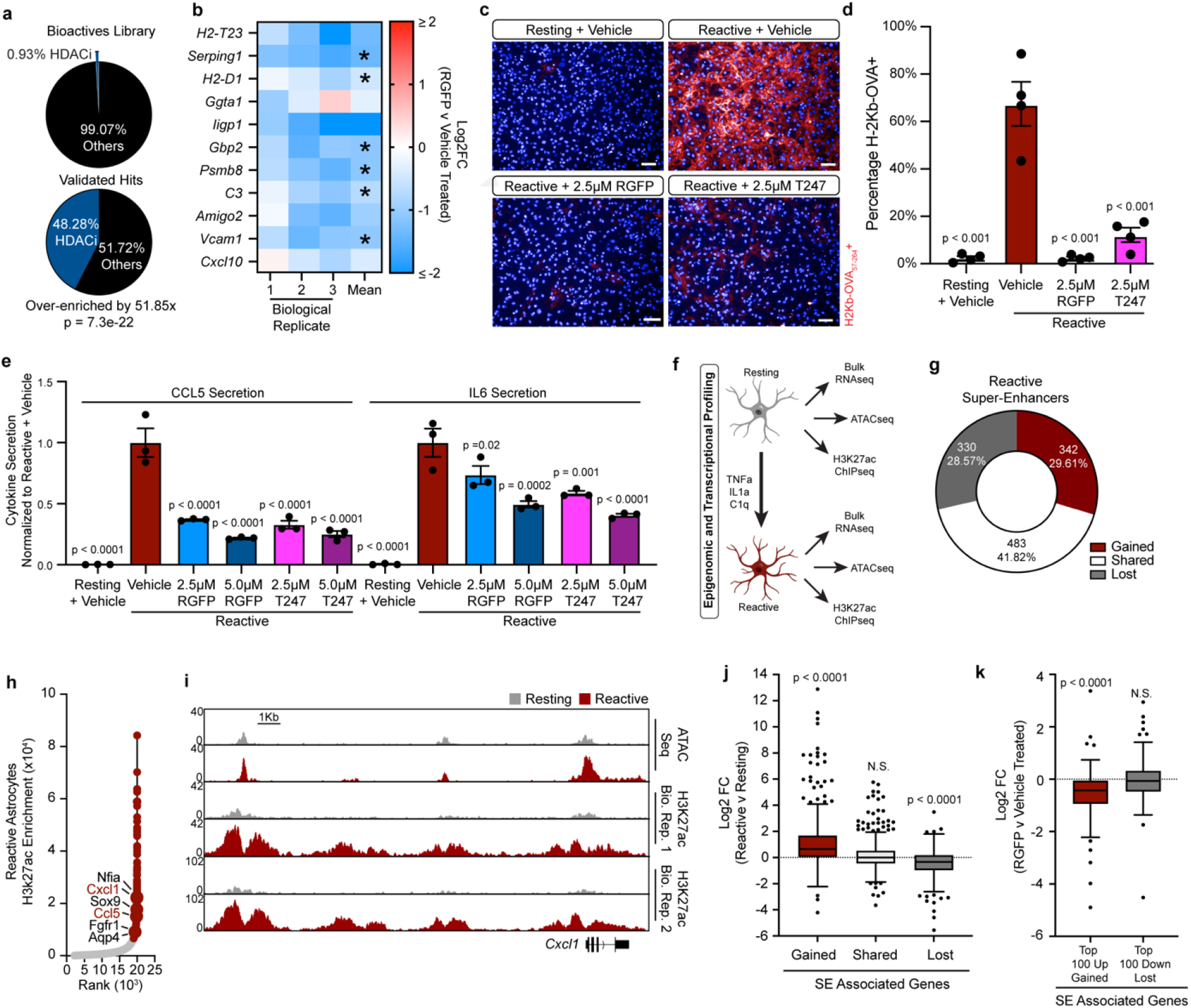
Chemical inhibition of HDAC3 modulates pathological reactive astrocytes. **a**, Proportion of HDAC inhibitor compounds enriched in primary screen validated hit list compared to the primary screen chemical library as a whole, showing that HDAC inhibitors are significantly enriched in the validated hit list. p-value generated by a hypergeometric test. **b**, Heatmap of the Log2 fold-change (Log2FC) between RGFP966 (RGFP) and vehicle (DMSO) treated pathological reactive astrocytes. Red is upregulated and blue is downregulated in RGFP treated cells. Data are presented as Log2FC for *n* = 3 biological replicates (independent astrocyte isolations) with mean and asterisks denoting a p < 0.05 calculated by DESeq2. **c**, Representative images of resting and reactive astrocytes treated with vehicle or the HDAC3 inhibitors RGFP or T247, then exposed to the OVA_257-264_ peptide and stained for MHC Class I bound to OVA_257-264_ (H-2Kb+OVA) in red. Scale bar is 50μm. **d**, Quantification of experiments represented in c. Data presented as mean ± s.e.m. *n* = 4 biological replicates (independent astrocyte isolations). p-values generated by a one-way ANOVA with Dunnett post-test for multiple comparisons to reactive plus vehicle control. e) Quantification of CCL5 and IL6 ELISAs performed on astrocyte conditioned media. Data presented as mean ± s.e.m for an *n* = 3 biological replicates (independent astrocyte isolations). p-value generated by a one-way ANOVA with Dunnett post-test for multiple comparisons to reactive plus vehicle control. **f**, Diagram of transcriptional and epigenetic data captured to analyze chromatin changes between resting and pathological reactive astrocytes. **g**, Distribution of gained, shared, and lost super-enhancers in pathological reactive astrocytes. **h**, Hockey-stick plot showing H3K27ac enrichment at enhancers in pathological reactive astrocytes. The enhancers are ranked and those in red were called as super-enhancers by ROSE analysis. Closest genes to each super-enhancer were called with HOMER, genes in red are targeted by gained super-enhancers in pathological reactive astrocytes while those in black are targeted by shared super-enhancers that were called in both resting and pathological reactive astrocytes. **i**, Example browser track for the gained super-enhancer gene *Cxcl1*. **j**, Tukey box and whisker plot showing the average Log2FC between pathological reactive and resting astrocytes for genes associated with gained, shared, and lost super-enhancers in pathological reactive astrocytes. p-value is generated with a one-sample Wilcoxon Signed Ranked test comparing to a hypothetical median of Log2FC = 0, which would designate no difference in expression between reactive and resting astrocytes. **k**, Tukey box and whisker plot showing the average Log2FC between RGFP and vehicle treated pathological reactive astrocytes for the top 100 upregulated genes associated with gained super-enhancers and the top 100 downregulated genes associated with lost super-enhancers in pathological reactive astrocytes. p-value is generated with a one-sample Wilcoxon Signed Ranked test comparing to a hypothetical median of Log2FC = 0 which would designate no difference in expression between reactive and resting astrocytes.

To understand how HDAC3 inhibitors regulate pathological reactive astrocytes, we sought to define the global chromatin and gene expression changes that occur as astrocytes transition to a pathological reactive state (Fig. 2f). H3K27ac ChIP-seq^24,25^ revealed substantial chromatin remodeling in pathological reactive astrocytes, including changes in super-enhancers that are regions of densely-packed active enhancers that control gene expression and cell state^26^. In pathological reactive astrocytes, nearly two-thirds of the super-enhancers were distinct from resting astrocytes demonstrating a substantial shift in the chromatin state (Fig. 2g). Expression of genes associated with gained super-enhancers in reactive astrocytes were significantly upregulated and included cytokines and chemokines involved in innate and adaptive immunity (*Cxcl1, Ccl5, Cxcl12*), while expression of genes associated with shared super-enhancers did not change and included astrocyte defining lineage markers (*Nfia. Sox9, Fgfr1*, and *Aqp4*)^27^ (Fig. 2h-j). HDAC3 inhibition by RGFP966 significantly decreased expression of gained super-enhancer genes but did not alter shared or lost super enhancer gene expression in reactive astrocytes (Fig. 2k). These data show that HDAC3 inhibition suppresses the transcripts induced in pathological reactive astrocytes by their dramatically altered chromatin landscape.

To uncover the regulators of astrocyte reactivity, we performed ATAC-seq and mined for motifs under open chromatin regions within gained super-enhancers in pathological reactive astrocytes. This analysis identified the RelA/p65 subunit of NF-κB as the top putative driver shaping the reactive astrocyte chromatin landscape (Extended Data Fig. 4a). Gene ontology analysis also highlighted NF-κB signaling as an enriched pathway in genes targeted by pathological reactive astrocyte gained super-enhancers (Extended Data Fig. 4b). Consistent with these data, we found a significant increase in nuclear RelA/p65 protein in reactive astrocytes that was abrogated by HDAC3 inhibition with RGFP966 (Extended Data Fig. 4c-e). Strikingly, we found that 82.75% (24/29) of the validated hits from our primary screen, including RGFP966, significantly inhibited RelA/p65 transcriptional activity in an orthogonal reporter assay (Extended Data Fig. 4f,g). Collectively, these data highlight a critical role of RelA/p65 in regulating pathological astrocyte reactivity and show that multiple molecular nodes, including HDAC3, can be targeted pharmacologically to impede RelA/p65 and modulate pathological reactive astrocytes.

### HDAC3 inhibition mediates a switch between pro- and anti-inflammatory astrocyte programs

To explore the role of RelA/p65 as a driver of reactive astrocytes in a pathological state, we performed RelA/p65 ChIP-seq in resting and pathological reactive astrocytes. We found that RelA/p65 DNA binding increased significantly as astrocytes become reactive (Fig. 3a), and that the majority of RelA/p65 direct target genes (genes with a RelA/p65 peak within 5Kb up- or down-stream of the transcription start site) had increased expression (60.43%, 113/187) (Fig. 3b). These included inflammatory cytokines and chemokines (*Cxcl10* and *Ccl2*), genes involved in MHC Class I antigen presentation (*H2-D1, B2m*, and *H2-K1*), and genes that define pathological reactive astrocytes (*C3* and *Gbp2*), (Fig. 3b). However, direct targets of RelA/p65 accounted for a small proportion (113/1839 upregulated transcripts, 9/1789 downregulated transcripts) of the overall gene expression changes in pathological reactive astrocytes, indicating that multiple regulatory programs may be mediating the generation of reactive astrocytes.

**Figure 3.**
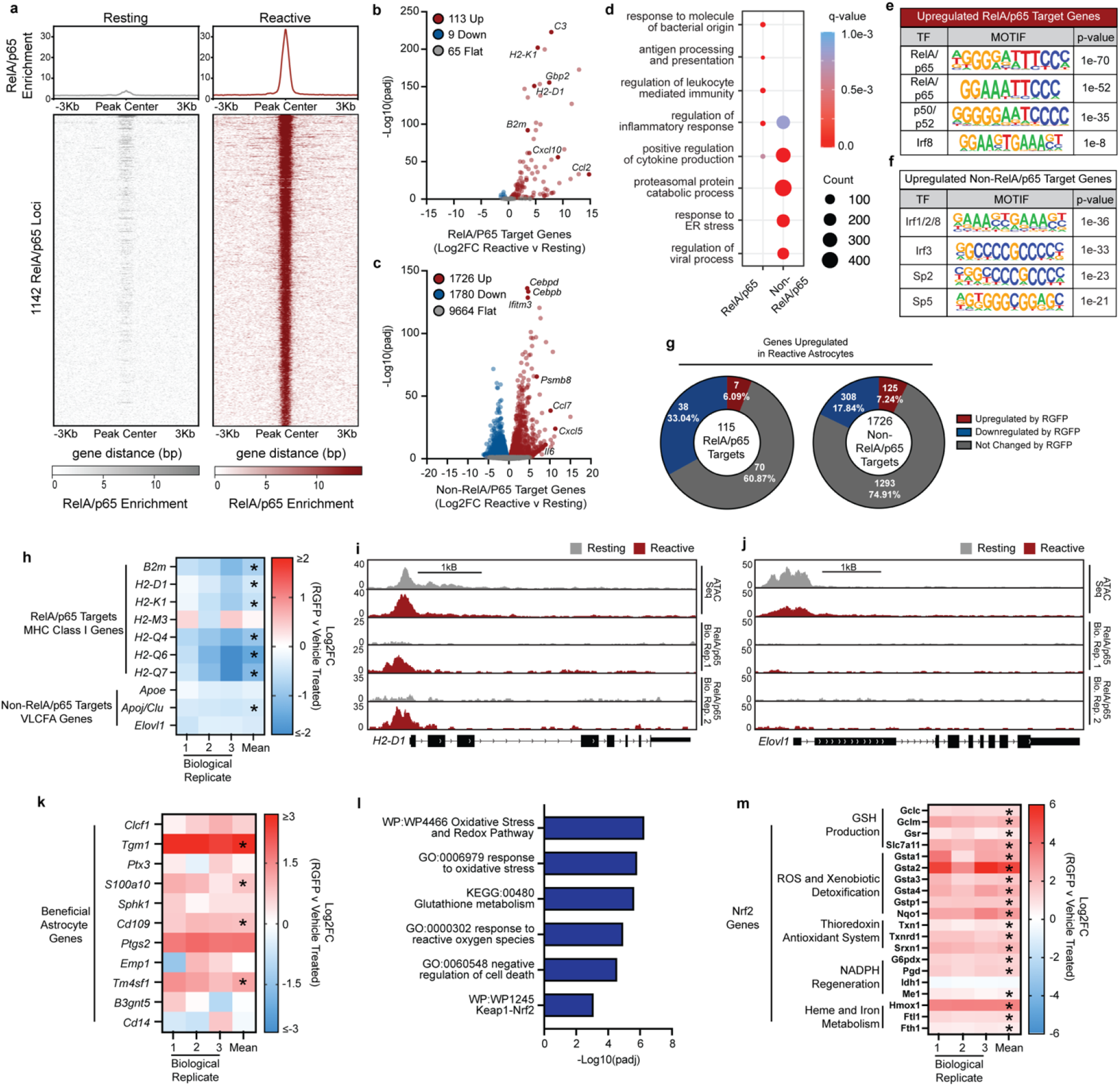
HDAC3 mediates a switch between pro-inflammatory and anti-inflammatory gene expression. **a**, RelA/p65 DNA binding aggregate heatmap and profile showing enriched RelA/p65 DNA binding at in reactive astrocytes compared to resting astrocytes. **b**, Volcano plot of RelA/p65 target genes (genes with a RelA/p65 peak within 5Kb up or downstream of the TSS). Of the 187 total RelA/p65 target genes in reactive astrocytes 60.43% (113/187) are upregulated while 4.81% (9/187) are downregulated and 34.76% (65/187) do not change. Log2FC and padj values were generated from bulk RNAseq analysis with DESEQ2 and with an *n* = 3 biological replicates (independent astrocyte isolations) per group. **c**, Volcano plot of the remaining Non-RelA/p65 target genes (genes without a RelA/p65 peak within 5Kb up or downstream of the transcription start site). Of the 14,364 Non-RelA/p65 target genes 13.10% (1726/13170) are upregulated in reactive astrocytes while 13.51% (1780/13170) are downregulated and 73.39% (9664/13170) do not change. **d**, Comparison of gene ontology terms called for RelA/p65 and Non-RelA/p65 target genes that are significantly upregulated in pathological reactive astrocytes. **e**, Transcription factor motifs identified with ATAC-seq at the promoter of the RelA/p65 target genes that are upregulated in reactive astrocytes compared to resting. **f**, Transcription factor motifs identified with ATAC-seq at the promoter of the Non-RelA/p65 target genes that are upregulated in reactive astrocytes compared to resting. **g**, Distribution of the effect that treatment with the HDAC3 specific inhibitor RGFP966 (RGFP) has on RelA/p65 and Non-RelA/p65 target genes that are upregulated in reactive compared to resting astrocytes. **h**, Heatmap showing the expression of the RelA/p65 MHC Class I genes and the non-RelA/p65 genes involved in very long-chain fatty acid synthesis and transport. Expression is shown as the Log2 fold-change (Log2FC) between 5uM RGFP966 and vehicle (DMSO) treated cells. Asterisks denote significance as called by DESeq2. **i**, Example browser track of the RelA/p65 target gene *H2-D1* from the MHC Class I gene set. **j**, Example browser track of the non-RelA/p65 target gene *Elovl1* that catalyzes the first and rate-limiting step in very long-chain fatty acid elongation. **k**, Heatmap showing the difference in expression between 5uM RGFP and vehicle-treated pathological reactive astrocytes for genes associated with a beneficial reactive astrocyte. Expression is shown as Log2FC for RGFP966 versus vehicle with asterisks denoting significance as called by DESeq2. **l**, Gene ontology analysis of genes significantly increased by RGFP treatment. **m**, Heatmap showing the difference in expression between 5uM RGFP and vehicle-treated pathological reactive astrocytes for cytoprotective Nrf2 target genes. Expression is shown as Log2FC for RGFP966 versus vehicle with asterisks denoting significance as called by DESeq2.

Non-RelA/p65 direct target genes with increased expression in pathological reactive astrocytes also included inflammatory cytokines and chemokines (*Cxcl5, Ccl7*, and *Il6*), transcription factors involved in acute inflammation (*Cebpb* and *Cebpd*), the immunoproteasome (*Psmb8, Psmb9*, and *Psmb10)*, interferon-induced genes (*Ifi47, Irf7, Iigbp1, Iigbp7*, and *Ifitm3*), and genes involved in lipid metabolism and transport (*Apoj/Clu* and *Elovl1*) (Fig. 3c). Gene ontology analysis of RelA/p65 and non-RelA/p65 direct target genes identified similar but not identical biological processes regulated by the two gene programs (Fig. 3d). However, transcription factor motifs enriched in the promoters of direct RelA/p65 and non-RelA/p65 direct target genes were distinct suggesting they are controlled by different mechanisms (Fig. 3e,f). HDAC3 inhibition by RGFP966 decreased expression of both RelA/p65 and non-RelA/p65 direct target genes in pathological reactive astrocytes (Fig. 3g). These included RelA/p65 direct target MHC Class I genes involved in the antigen presentation function blocked by RGFP966, and non-RelA/p65 direct target genes associated with the metabolism and transport of very-long chain fatty acids that were recently shown to partially mediate pathological reactive astrocyte toxicity to neurons and oligodendrocytes^6^ (Fig. 3h-j). These data show that HDAC3 inhibition broadly suppresses the multiple transcriptional programs that regulate pathological reactive astrocyte formation.

While most transcripts changed by RGF9666 in reactive astrocytes were decreased, nearly 40% of all RGFP966 changed transcripts (539/1380) were increased. Transcripts increased in reactive astrocytes by RGFP966 treatment included numerous genes associated with a beneficial reactive astrocyte state^1^ (Fig. 3k). Moreover, gene ontology analysis of genes induced by RGFP966 treatment was enriched for terms associated with response to oxidative stress, glutathione metabolism, and KEAP1-NRF2 signaling (Fig. 3l). NRF2 activation in astrocytes has been shown to protect against neurodegeneration and negatively regulate inflammation^3,28^. Importantly, we found that HDAC3 inhibition with RGFP966 activated NRF2 signaling. Cytoprotective Nrf2 target genes^29^ including those involved in glutathione production, antioxidant systems, NADPH regeneration, and heme and iron metabolism were significantly induced by RGFP966 treatment (Fig. 3m). These findings show that HDAC3 inhibition not only mitigates proinflammatory target gene expression, but also promotes expression of genes associated with an anti-inflammatory beneficial astrocyte state. This suggests that HDAC3 acts as a molecular switch between reactive astrocyte states where HDAC3 inhibition promotes an anti-inflammatory and protective astrocyte phenotype.

### Pharmacological inhibition of HDAC3 promotes neuroprotection *in vivo*

To examine the therapeutic effect of modulating reactive astrocytes, we tested whether chemical HDAC3 inhibition could reduce pathological reactive astrocytes *in vivo*. With daily intraperitoneal (i.p.) dosing of 10mg/kg, RGFP966 crossed the blood-brain barrier and reached a concentration in brain tissue near *in vitro* IC50 levels required to inhibit astrocyte reactivity (Extended Data Fig. 5a). RGFP966 increased gross acetylated histone 4 (AcH4) brain levels *in vivo* demonstrating a pharmacodynamic effect to inhibit the histone deacetylase activity of HDAC3 (Extended Data Fig. 5b-d). We used the well-established model of systemic LPS-induced neuroinflammation^30,31^ to test whether HDAC3 inhibition with RGFP966 could modulate pathological reactive astrocytes *in vivo*. Compared to vehicle, treatment with 10mg/kg RGFP966 successfully decreased the formation of *Gbp2*-positive pathological reactive astrocytes in LPS-exposed mice (Fig. 4a-c). Importantly, RGFP966 treatment had no effect on general astrocyte reactivity as measured with GFAP and had no effect on microglial activation measured with IBA-1(Extended Data Fig. 5e-k). This shows that HDAC3 inhibition with RGFP966 can specifically decrease the formation of the pathological reactive astrocyte state *in vivo*.

**Figure 4.**
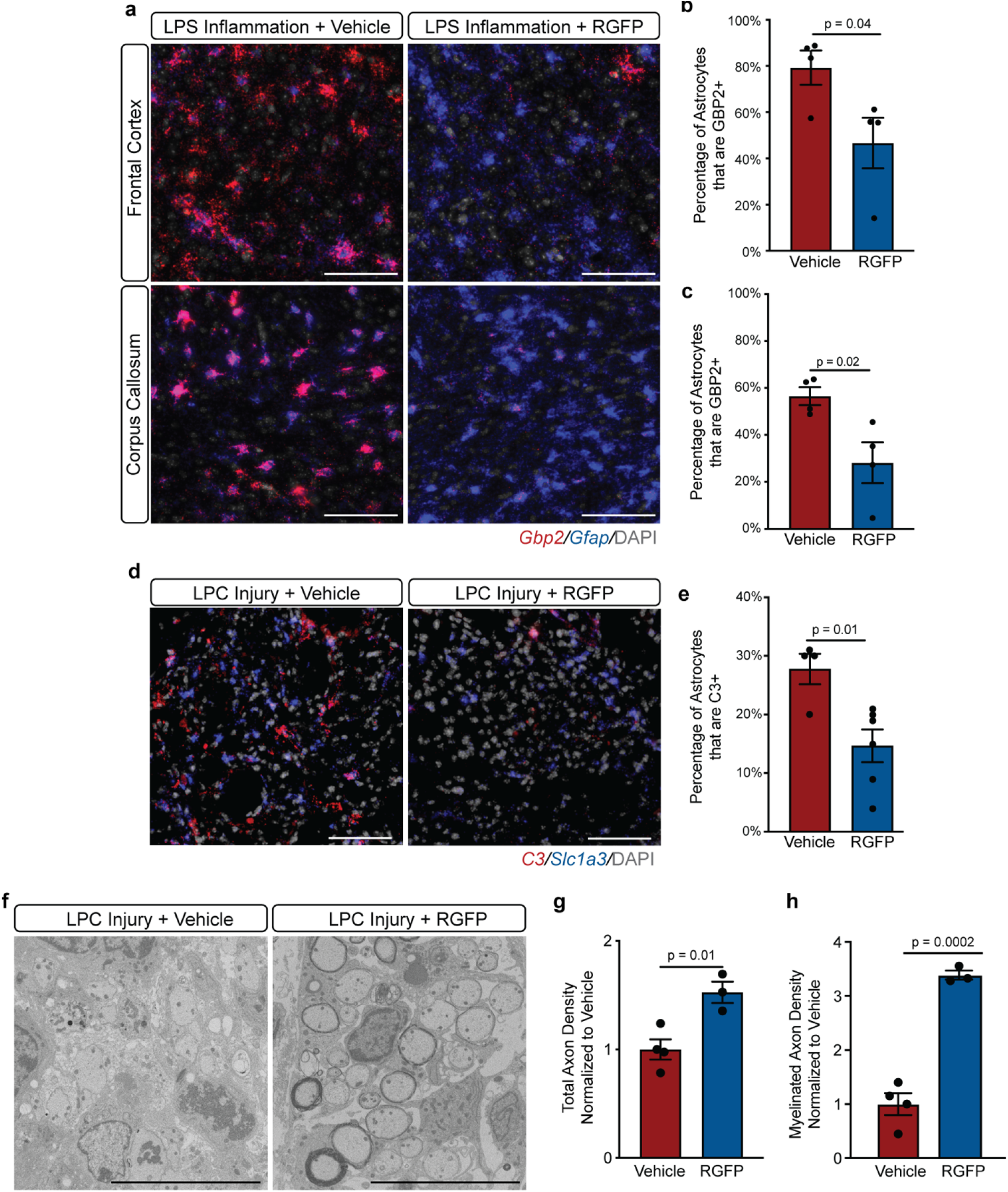
HDAC3 inhibition blocks reactive astrocyte formation *in vivo* and promotes axonal protection. a) Representative images of *Gbp2* (red) and *Gfap* (blue) *in situ* hybridization from the frontal cortex and corpus callosum of mice after 48hrs of systemic lipopolysaccharide (LPS) exposure and treated with vehicle or 10mg/kg RGFP. Scale bar is 100um. **b-c**, Quantification of *in situ* data represented in a. The percentage of astrocytes that are GBP2+ in the **b**, frontal cortex or **c**, corpus callosum of mice exposed to LPS and treated with vehicle or 10mg/kg RGFP is presented as mean ± s.e.m for *n* = 4 biological replicates (2 male and 2 female mice) with p-value generated by Student’s unpaired two-tailed t-test. **d**, Representative images of *C3* and *Slc1a3 in situ* hybridization in the dorsal column of lysolecithin (LPC) lesioned mice at 12 days post lesion and treated with vehicle or 10mg/kg RGFP. Scale bar is 50um. **e**, Quantification of *in situ* data represented in D. The percent of astrocytes that are *C3* positive in LPC-exposed mice treated chronically with vehicle 10mg/kg RGFP. Data are presented as the mean ± s.e.m for *n* = 4-6 biological replicates (mice) with p-value generated by Student’s unpaired two-tailed t-test. **f**, Representative electron microscopy (EM) images of LPC lesioned mice treated with vehicle or 10mg/kg RGFP. Scale bar is 10um. **g**, Quantification of axon density in EM images represented in f. Total axon density normalized to control for LPC lesioned mice treated with vehicle (*n* = 4 mice) or 10mg/kg RGFP (*n* = 3 mice) is presented as mean ± s.e.m. with p-value generated by Student’s unpaired two-tailed t-test. h) Quantification of remyelinated axon density in EM images represented in f. Myelinated axon density normalized to control for LPC lesioned mice treated with vehicle or 10mg/kg RGFP is presented as mean ± s.e.m for *n* = 3-4 biological replicates (mice) with p-value generated by Student’s unpaired two-tailed t-test.

Finally, we examined whether decreasing pathological reactive astrocyte formation by pharmacologically inhibiting HDAC3 could promote neural protection. To do this, we used a toxin-based model of CNS tissue damage where lysolecithin (lysophosphatidylcholine; LPC) is injected into the dorsal column of the spinal cord, leading to myelinating oligodendrocyte loss, astrogliosis, and both acute and continued axonal degeneration^32,33^. We first confirmed that LPC-induced injury leads to the formation of *C3*-positive pathological reactive astrocytes, that RGFP966 significantly decreased the formation of these *C3*-positive pathological reactive astrocytes, and that RGFP966 had no effect on general microgliosis or astrogliosis in this injury model (Fig. 4d,e; Extended Data Fig. 6a-d). Using electron microscopy to assess tissue pathology, we compared spinal cord sections from RGFP966-treated versus vehicle-treated LPC mice and found that HDAC3 inhibition with RGFP966 promoted axonal protection (Fig. 4f). At 12 days post-LPC injury, RGFP966 treatment significantly increased the density of total axons and the density of myelinated axons (Fig. 4g,h). These data show that pathological reactive astrocytes play a role in the loss of both unmyelinated and myelinated axons in the LPC injury model and that pharmacological inhibition of HDAC3 suppressed pathological reactive astrocytes and promoted axonal protection following *in vivo* tissue damage.

## Conclusions

Here we show the development of an astrocyte discovery platform to identify modulators of reactive astrocytes. A phenotypic chemical screen identified HDAC3 as a functional regulator of a pathological reactive astrocyte state change. We demonstrate that pathological reactive astrocytes represent a cell state change that involves global reorganization of the chromatin landscape driven in part by RelA/p65. HDAC3 inhibition in reactive astrocytes suppresses RelA/p65 and non-RelA/p65 proinflammatory gene programs, while increasing expression of anti-inflammatory and beneficial reactive astrocyte associated genes. We further show that inhibition of HDAC3 decreases the formation of pathological reactive astrocytes in an *in vivo* model of neuroinflammation. These findings support the role of HDAC3 as a molecular switch to control astrocyte polarization between pathological or beneficial cell states and suggest that therapeutics targeting HDAC3 in neurodegenerative disease may have the dual benefit of blocking pathological reactive astrocyte formation while promoting the formation of beneficial reactive astrocytes to protect neurons and promote repair. We support this by showing in a model of toxin-induced CNS damage that pharmacological inhibition of HDAC3 decreases the formation of pathological reactive astrocytes and promotes neural protection. These findings provide a foundation for identification of additional pathological reactive astrocyte modulating chemicals, specify a deeper understanding of the molecular regulators of pathological reactive astrocyte gene expression, and support the development of reactive astrocyte-targeted therapies for neurodegenerative diseases.

## Methods

### Mouse studies

All primary cell isolation and LPS studies were performed at Case Western Reserve University. All LPC studies were performed at George Washington University Medical Center. All animal procedures were in accordance with the National Institutes of Health Guidelines for the Care and Use of Laboratory Animals and were approved by the Case Western Reserve University Institutional Animal Care and Use Committee or the George Washington University Medical Center Institutional Animal Care and Use Committee.

### Lipopolysaccharide (LPS) model of neuroinflammation

C57BL/6N mice were purchased from Charles River (Wilmington, MA). Male and female mice at 7 weeks of age were injected i.p. with either RGFP966 vehicle (30% hydroxypropyl-β-cyclodextrin, 0.1M sodium acetate, and 10% DMSO) or 10mg/kg RGFP966 daily for 11 days. After which mice were injected i.p. daily for two days with either LPS vehicle (saline) plus RGFP966 vehicle, 5mg/kg LPS plus RGFP966 vehicle, or 5mg/kg LPS plus 10mg/kg RGFP966. Animals were then sacrificed and processed for immunohistochemistry.

### Lysolecithin (LPC) injection model of focal tissue damage

Focal tissue damage in the spinal cord was induced by the injection of 1% LPC solution. 10–12-week-old C57BL/6 female mice were anesthetized using isoflurane and a T10 laminectomy was performed. 1μL of 1% LPC was infused into the dorsal column at a rate of 15 mL/hour. The animals were euthanized at day twelve after the laminectomy (n= 3-4 per group). Animals received either RGFP966 vehicle or 10mg/kg RGFP966 daily by i.p. injections that began 1 day prior to LPC injection and ended 11 days after LPC injection. Experiments were terminated 12 days after LPC injection and tissue was then processed for immunohistochemistry or electron microscopy.

### Isolation and generation of primary astrocytes

Timed-pregnant C57BL/6N mice were purchased from Charles River (Wilmington, MA). Brains from mice of the C57BL/6N strain were extracted at postnatal day 2 (P2). A gross dissection of these brains was performed to isolate the cortices which were then dissociated according to manufacturer’s instructions using the Miltenyi Tumor Dissociation Kit (Miltenyi, 130-095-929). After dissociation, cells were plated in flasks coated with poly-L-ornithine (Sigma, P3655) and laminin (Sigma, L2020). Cells were cultured for 24 hours in media consisting of Dulbecco’s Modified Eagle Medium/Nutrient Mixture F-12 (DMEM/F-12; ThermoFisher 11320033), N-2 MAX Supplement (ThermoFisher, 17502048), B-27 Supplement (ThermoFisher, 17504044), GlutaMAX Supplement (ThermoFisher, 35050079), Penicillin-Streptomycin (ThermoFisher, 15070063), and 20ng/mL FGF-2. After 24 hours, the cells were switched to astrocyte enrichment media, comprised of 1:1 DMEM (ThermoFisher, 11960044) and Neurobasal Medium (ThermoFisher, 21103049) supplemented with GlutaMAX (ThermoFisher, 35050061), Sodium Pyruvate (ThermoFisher, 11360070), N-2 MAX Supplement (R&D, AR009), 5ug/ml N-acetyl cysteine (Sigma, A8199), Penicillin-Streptomycin (ThermoFisher, 15070-063), 5 ng/mL HB-EGF (R&D Systems, 259-HE-050), 10 ng/mL CNTF (R&D Systems, 557-NT-010), 10 ng/mL BMP4 (R&D Systems, 314-BP-050), and 20 ng/mL FGF2 (R&D Systems, 233-FB-01M) to proliferate. Media was changed every 48 hours. Once confluent, astrocytes were either cryopreserved or passaged once and then cryopreserved. To conduct a terminal experiment, cryopreserved astrocytes were removed from liquid nitrogen storage and thawed in astrocyte maturation media (1:1 DMEM and Neurobasal Medium with GlutaMAX Supplement, Sodium Pyruvate, N-2 MAX Supplement, N-acetyl cysteine supplemented with 5 ng/mL HB-EGF, 10 ng/mL CNTF, 50 ng/mL BMP4, and 20 ng/mL FGF-2) for 48 hours, followed by resting astrocyte media (1:1 DMEM and Neurobasal Medium with 5ng/ml HB-EGF) for another 72 hours. After this five-day thawing and maturation protocol, experimental treatments could be applied to these mature astrocytes in culture.

### Primary screen and secondary dose response screen

All liquid handling was performed using a BioTek EL406 Washer Dispenser. Astrocytes were thawed and plated, as described for terminal experiments above, onto 384-well plates as described above at a density of 500 cells/mm^2^. A Perkin Elmer Janus G3 Varispan Automated Workstation was then used to treat cells with small-molecules with one molecule per well at a concentration of 2uM in 384-well plates, followed one hour later by 3 ng/mL IL1α (Sigma #I3901), 400 ng/mL C1q (MyBioSource #MBS143105), and 30 ng/mL TNF (R&D Systems #210-TA-020). After incubation for 24 hours, the cells were fixed using 4% paraformaldehyde and stained for GBP2 (Proteintech #11854-1-AP) using the procedure detailed in the immunocytochemistry section below, then imaged using the PerkinElmer Operetta CLS High-Content Analysis System. Images were analyzed using automated PerkinElmer Columbus Image Analysis Software. For analysis, toxic chemicals were first removed, a chemical was considered toxic if it decreased the counted number of live cells in the well by greater than 30% compared to reactive astrocyte plus DMSO vehicle control wells. Hits were then determined as those compounds that decreased the number of GBP2-positive reactive astrocytes by greater than or equal to 90% compared to reactive astrocyte plus DMSO vehicle control wells.

Secondary dose curve screens were performed exactly as described for the primary screen except with custom generated dose curve plates containing the following drug concentrations: 6uM, 3uM, 1.5uM, 0.75uM, 0.37uM, 0.18uM, 0.09uM.

### Z-Score Calculation

Z-score is a standard measure of drug screen quality. To calculate the Z’ standard where σ = standard deviation and μ = mean of either the positive or negative control wells:

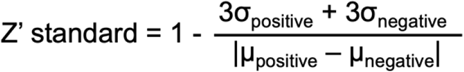

To calculate Z’ Robust where (mad) = mean absolute deviation and 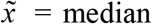 of either the positive or negative control wells:

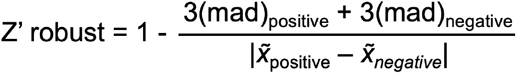

### Generation of *Hdac3* knockout astrocytes

To generate *Hdac3* knockout astrocytes cells were isolated from *Aldh1l1-CreERT*^*2*^*;Hdac3*^*fl/fl*^ mice, generated by crossing *Aldh1l1-CreERT*^*2 22*^ (Jax #031008) and *Hdac3*^*fl/fl*^ (Jax #024119)^23^ mice. *Aldh1l1-CreERT*^*2*^*;Hdac3*^*fl/fl*^ astrocytes were thawed and matured as described above then treated with 1uM tamoxifen (Selleck, S7827) for three days followed by 4-5 days to allow recombination. Cells were then treated with 3 ng/mL IL1α (Sigma #I3901), 400 ng/mL C1q (MyBioSource #MBS143105), and 30 ng/mL TNF (R&D Systems #210-TA-020) and 24hrs later fixed and stained as described in immunocytochemistry section below.

### Human astrocyte generation

Human induced pluripotent stem cells (iPSCs) were cultured to differentiate into astrocytes as previously described^34^. In short, iPSC colonies were placed in neural induction media for 10 days until neural rosettes could be picked, dissociated, and plated in a glial expansion medium. These cells were allowed to proliferate and become a homogenous population of glial progenitor cells (GPCs) over eight passages on poly-L-ornithine and laminin-coated plates. These GPCs were then passaged onto a Matrigel-coated plate to culture in an astrocyte induction media for two weeks, which was followed by culturing for another four weeks in an astrocyte maturation medium.

### Real-time polymerase chain reaction (qPCR)

Cells were lysed in TRIzol and total RNA was extracted with phenol-chloroform followed by spin columns from the RNEasy Mini Kit (Qiagen, 74104). RNA quality and quantity was determined using a NanoDrop spectrophotometer. The RNA was then reverse transcribed using the iScript cDNA synthesis kit (BioRad, 1708891) according to the manufacturer’s instructions. Real-time qPCR was performed using the Taqman Gene Expression Master Mix (Applied Biosystems, 4369016) and the Taqman assay probes for human: *GBP2* (Thermo Fisher Assay ID: Hs00894837_m1) and *PSMB8* (Hs00544758_m1).

### ViewRNA ISH *in vitro*

*In situ* hybridization of cells in culture was performed with ThermoFisher ViewRNA according to manufacturer’s instructions without modification (ThermoFisher #QVC0001). Briefly, cells were fixed and permeabilized followed by hybridization with target-specific probes for 2hrs at 40C. Target probe signal was then amplified and signal was imaged using the PerkinElmer Operetta CLS High-Content Analysis System and analyzed using automated scripts in PerkinElmer Columbus Image Analysis software. Probes for *C3* (ThermoFisher, VB4-3112231-VC) and *Serping1* (ThermoFisher, VB4-3114232-VC).

### Enzyme-linked immunosorbent assay (ELISA) and cytokine profiling

Secreted cytokine levels were measured by ELISA and cytokine profiling arrays. ELISA assays were performed according to the manufacturer’s instructions for CCL5 (R&D Systems, DY478-05) and IL6 (DY406-05). Relative CCL5 and IL6 concentrations were measured based on absorbance measured by Synergy Neo2 (BioTek) plate reader. Secreted cytokines were measured using the Proteome Profiler Array Mouse XL Cytokine Array Kit (R&D Systems, ARY028) according to manufacturer’s instructions.

### OVA_257-264_ peptide antigen presentation assay

Mouse primary astrocytes were cultured and treated with reactive astrocyte driving cytokines with or without the HDAC3 specific inhibitors, RGFP966 and T247. After 24 hours of treatment, these cells were cultured in the presence of the OVA_257-264_ peptide (GenScript RP10611 or Sigma S7951). After 12 hours, the cells were live-stained for two hours using a conjugated antibody targeted against an OVA_257-264_ peptide antigen (BioLegend, 141605). The cells were then fixed with 4% paraformaldehyde and imaged using the PerkinElmer Operetta CLS High-Content Analysis System. Images were analyzed using the automated Columbus Image Data Storage and Analysis System to identify the extent to which HDAC3 inhibitors decreased OVA_257-264_ antigen presentation.

### NF-κB Jurkat reporter assay

Jurkat NF-κB reporter cells were purchased from BPS Bioscience (60651). Cells were grown in manufacturer recommended growth media of RPMI 1640 with 10% FBS, 1x Pen/Strep, and 1mg/mL Geneticin. To test whether drugs inhibit NF-κB activity, the reporter cells were exposed to drugs or DMSO vehicle for one hour before the addition of 3 ng/mL IL-1a (Sigma, I3901), 400 ng/mL C1q (MyBioSource, MBS143105), and 30 ng/mL TNF (R&D Systems, 210-TA-020). Cells were then incubated for 24hrs, then luciferase activity was measured with the One-Step Luciferase Assay System (BPS Bioscience, 60690-1) according to manufacturer’s instructions with a Synergy Neo2 plate reader (BioTek).

### Immunocytochemistry

For immunocytochemistry, cells were fixed with ice-cold 4% PFA for 15 minutes at room temperature, washed three times with PBS, blocked and permeabilized with 10% donkey serum and 0.1% Triton X-100 in PBS for 1hr, and then stained with primary antibodies overnight followed by three washes with PBS, and then 1hr incubation with Alexa fluor secondary antibodies and DAPI. Stained cells were imaged using the PerkinElmer Operetta CLS High-Content Analysis System and analyzed using automated scripts in PerkinElmer Columbus Image Analysis software. Primary antibodies used were GBP2 (ProteinTech, 11854-1-AP), Vimentin (BioLegend, 919101), AQP4 (Sigma, HPA014784), GLT-1 (Novus Biologicals, NBP1-20136), ALDH1L1 (Novus Biologicals, NBP2-50045), and HDAC3 (BioLegend, 685202).

### Single-cell RNAseq sample preparation and analysis

Resting and reactive astrocytes were lifted from culture plates using TrypLE (ThermoFisher, 12563011) and collected in 1x PBS with 1% bovine serum albumin (BSA). Cells were then spun down at 1000rpm for 10 minutes. After which cells were washed once with 1x PBS with 1% BSA before being filtered through 40um FlowMi Tip Strainers (VWR, 10032-802). Cells were then diluted with 1x PBS with 1% BSA and loaded onto the Chromium 10x Controller according to manufacturer’s instructions. Following partitioning of single-cells in gel bead emulsions, reverse-transcriptions and library preparation were performed according to 10x Single Cell 3’ v2 chemistry kit instructions (v2 kit since discontinued). Finally, libraries were sequenced by the Case Western Reserve University Genomics Core on an Illumina HiSeq2500 with paired-end 50bp reads and a target sequence depth of 50,000 reads per cell. Sequence data were first processed by 10x Cell Ranger v3.1 using default settings to generate a gene expression matrix and then all downstream analysis was performed with the R package, Seurat v4.0^12^.

### Bulk RNAseq sample preparation and analysis

Total RNA was extracted from resting and reactive astrocytes using the same procedure as described for qPCR and sent to Novogene for mRNA Sequencing. For gene expression analysis, reads were mapped to the mm10 genome using kallisto v0.46.1 (https://pachterlab.github.io/kallisto/)^13^. Transcripts were summarized to the gene level with tximport v1.2 (https://bioconductor.org/packages/release/bioc/html/tximport.html)^14^. Normalized expression and differential gene expression were then generated using DESeq2 v1.32.0 (https://bioconductor.org/packages/release/bioc/html/DESeq2.html)^15^.

### H3K27ac and RelA/P65 Chromatin immunoprecipitation sequencing (ChIP-seq)

Nuclei isolation and chromatin shearing were performed using the Covaris TruChIP protocol following manufacturer’s instructions for the “high-cell” format. In brief, 5 million (H3K27Ac) or 20 million resting and reactive (RelA/P65) were cross-linked in ‘‘Fixing buffer A’’ supplemented with 1% fresh formaldehyde for 10 minutes at room temperature with oscillation then quenched for 5 minutes with ‘‘Quench buffer E.’’ These cells were then washed with PBS and either snap frozen and stored at -80C or immediately used for nuclei extraction and shearing per the manufacturer protocol. The samples were sonicated with the Covaris S2 using the following settings: 5% Duty factor 4 intensity for four 60 seconds cycles. Sheared chromatin was cleared and incubated overnight at 4C with primary antibodies that were pre-incubated with protein G magnetic DynaBeads (Thermo Fisher, 10004D). Primary antibodies used included anti-H3K27Ac (Abcam, ab4729) and anti-RelA/P65 (Cell Signaling Technology, 8242). These beads were then washed, eluted, reverse cross-linked, and treated with Rnase A followed by proteinase K. ChIP DNA was purified using Ampure XP beads (Aline Biosciences, C-1003-5) and then used to prepare Illumina sequencing libraries as described previously^16^. Libraries were sequenced on the Illumina HiSeq2500 with single-end 50bp reads with a read-depth of at least 20 million reads per sample.

For peak calling, reads were quality and adaptor trimmed using Trim Galore! Version 0.4.1. Trimmed reads were aligned to mm10 with Bowtie2 version 2.3.2^17^. Duplicate reads (potential artifacts of PCR in library preparation) were removed using Picard MarkDuplicates. Peaks were called with MACS version 2.1.1^18^. Peaks were visualized with the Integrative Genomics Viewer (IGV, Broad Institute). Peaks were compared and contrasted using bedtools implemented in the R software environment using the BedtoolsR package (http://phanstiel-lab.med.unc.edu/bedtoolsr.html) and peaks were assigned to the nearest gene using the R packages ChIPSeeker^19^, ChIPpeakanno^20^, and GenomicRanges^21^. Super-enhancers were called using rank ordering of super-enhancers (ROSE) analysis package^24,25^. Super-enhancers and genes targeted by super-enhancers were generated for the H3K27ac biological replicate with the strongest signal and key findings confirmed with a second biological replicate.

### Omni Assay for Transposase-Accessible Chromatin using sequencing (ATAC-seq)

Omni ATAC-Seq was performed on 50,000 resting and reactive astrocytes following the protocol outlined in Corces et al.^22^ In brief, nuclei were extracted from cells and treated with transposition mixture containing Nextera Tn5 Transposase for (Illumina, FC-121-1030). Transposed fragments were then purified using QIAGEN MinElute columns (QIAGEN, 28004), PCR amplified, and libraries were purified with PCRClean DX purification system (Aline Biosciences, C-1003-5) with a sample to bead ratio of 1:1.2. Final libraries were sequenced on the illumina HiSeq2500 with single-end 50bp reads and nearly 100 million reads per sample. Reads were aligned to the mm10 mouse genome following the same pipeline used for ChIP-seq data (see ChIP-seq and analysis) and peaks were called using the “narrowPeaks” function of MACS version 2.1.1, as outlined for RelA/P65 ChIP-seq (see ChIP-seq and analysis).

### Motif enrichment analysis

Motifs were called under RelA/P65 peaks, ATAC-seq peaks, or regions of gained or lost H3K27Ac using HOMERv4.11.1 (Heinz et al., 2010). The findMotifsGenome.pl tool was used with mm10 as the reference genome.

### Pharmacokinetics

C57BL/6 adult mice were injected i.p. with 10mg/kg RGFP966 daily for 11 days. Mice were then perfused with saline to remove blood from the brain. Brain tissue was then collected and snap frozen. Brain tissues were thawed at room temperature and homogenized in PBS. Calibration standards and study samples were extracted with 3x volume of acetonitrile containing 0.1% formic acid with 200 ng/ml internal standard solution. Samples were then each vortexed for 1 minute, then transferred to an Eppendorf R5417R and centrifuged at 14000 rpm for 7 minutes. Following extraction of tissue, homogenate calibrators and study samples were transferred directly to an autosampler microtiter plate for analysis. Samples were analyzed by LC-MS-MS in the positive ion electrospray ionization mode.

### Immunohistochemistry

Mice were perfused with PBS followed by 4% paraformaldehyde, after which brains were extracted and cryopreserved in 30% sucrose, then frozen in OCT and sectioned. To stain, slides were washed with PBS and then incubated overnight with primary antibody. After primary incubation, slides were then washed and labeled with Alexa Fluor secondary antibodies (ThermoFisher). Images were captured on a Hamamatsu Nanozoomer S60 Digital slide scanner with NDPview 2.0 software. Image analysis was performed using automated scripts with Perkin Elmer Columbus Image Analysis Software. Primary antibodies used were AcH4 (EMD Millipore, 06-866), GFAP (DAKO, 685202), and IBA-1 (Abcam, 685202)

### Western Blotting

Protein samples were collected in RIPA buffer (Sigma, R0278) with Halt Protease and Phosphatase Inhibitor (ThermoFisher, 78441). Total protein concentration was determined by BCA assay (ThermoFisher, 23225). Equal amounts of total protein were loaded onto 4-12% Bis-Tris gels (Invitrogen). Proteins were then separated by gel-electrophoresis and transferred to PVDF membranes (ThermoFisher, LC2002). Blots were probed with RelA/p65 (Cell Signaling Technology, 8242) and beta-Actin (Sigma, A3854), developed using SuperSignal West Pico Plus Chemiluminescent Substrate (ThermoFisher, 34577), and visualized using a Li-Cor Odyssey XF Imager.

#### *In situ* hybridization with RNAscope

*In situ* hybridization for *in vivo* studies was performed using RNAscope Multiplex Fluorescence V2 Assay (ACD Bio, 323136) according to manufacturer’s instructions for fixed frozen samples. Briefly, tissue was prepared by first dehydrating with increasingly higher percentages of ethanol, then dried, blocked with hydrogen peroxide, followed by antigen retrieval for 5 minutes, dried again, and then protein was digested using provided Protease III. RNA-Targeting probes purchased from ACD Bio were then annealed at 40C for 2 hours followed by washing and a series of amplification steps before finally tagging the RNA with Opal Dye fluorophores (Perkin Elmer). *In situ* hybridization Images were captured on a Hamamatsu Nanozoomer S60 Digital Slide scanner with NDPview 2.0 software. Image analysis was performed using automate scrips with Perkin Elmer Columbus Image Analysis Software. The following mouse specific RNAscope probes were used: *Slc1a3* (ACD Bio, 430781), *Gbp2* (ACD Bio, 572491), *Gfap* (ACD Bio, 313211), and *C3* (ACD Bio, 417841).

### Electron microscopy sample preparation and analysis

Samples were processed as previously described^23^. Briefly, mice were perfused with 4% PFA, 2% gluteraldehyde, and 0.1 M sodium cacodylate. Spinal cords were extracted, and samples were osmicated, stained *en bloc* with uranyl acetate and embedded in Embed 812, an Epon-812 substitute (Electron Microscopy Sciences). Thin sections were cut, carbon-coated and imaged either on a Helios NanoLab 660 Scanning Electron Microscope (FEI)

### Gene ontology (GO) and gene set enrichment analysis (GSEA)

GO analysis was performed using gProfiler (https://biit.cs.ut.ee/gprofiler/gost)^24^ with a significance threshold set at false discovery rate (FDR) < 0.05 and calculated by Benjamini-Hochberg FDR. Redundant GO terms were removed using REVIGO^35^ (http://revigo.irb.hr/) using default settings. Comparisons between RelA/p65 direct target genes and non-RelA/p65 direct target genes GO terms was performed with the R package, clusterProfiler^36^ v4.0.5 (https://bioconductor.org/packages/release/bioc/html/clusterProfiler.html), following the tutorial for “Biological Theme Comparison” at https://yulab-smu.top/biomedical-knowledge-mining-book/index.html. GSEA scores were generated for curated gene sets from publicly-available data. GSEA was run using classical scoring, 1000 gene-set permutations, and signal-to-noise metrics. Normalized enrichment scores (NES) and FDR were calculated by the GSEA software^37^ (https://www.gsea-msigdb.org/gsea/index.jsp).

### Statistical analysis

Unless otherwise noted, GraphPad Prism was used to perform statistical analyses The statistical tests used and description of data presentation and sample size can be found in each figure legend. Unless otherwise noted, sample sizes were determined by reference to previous literature. For all Tukey box-and-whisker plots, the middle line of the box is the median, the box extends from the 25^th^ to 75^th^ percentile, the upper whisker is placed at the 75^th^ percentile plus 1.5x the inter-quartile distance, the lower whisker is placed at the 25^th^ percentile plus 1.5x the inter-quartile distance, and any individual data points that fall outside of the upper and lower whiskers are plotted.

## Data availability

All datasets generated in this study have been deposited in Gene Expression Omnibus (https://www.ncbi.nlm.nih.gov/geo/) under SuperSeries accession code GSE185215 with subseries for RNA-seq (GSE185212), scRNA-seq (GSE184437), ChIP-seq (GSE185606), and ATAC-seq (GSE185605).

## Acknowledgments

This study was supported by grants from the National Institutes of Health R35NS116842 (P.J.T.), F30HD096784 (K.C.A.), and T32GM007250 (K.C.A), and the Hartwell Foundation (B.L.L.C), as well as institutional support from the Case Western Reserve University School of Medicine and philanthropic support from the Fakhouri, R. Blane & Claudia Walter, Long, Goodman, Geller, and Weidenthal families, and the Research Institute for Children’s Health. Additional support was provided by the Small Molecule Drug Development and Genomics core facilities of the CWRU Comprehensive Cancer Center (P30CA043703), the CWRU Light Microscopy Imaging Center (S10-OD016164), and the University of Chicago Genomics Facility. The authors thank Drew Adams, Yuriy Federov, Bill Harte, Marissa Scavuzzo, Erin Cohn, Mayur Madhavan, and Matt Elitt for technical assistance and/or discussion.

## Author contributions

B.L.L.C. and P.J.T. conceived and managed the overall study. B.L.L.C. and J.D.K. performed, quantified, and analyzed all *in vitro* studies. H.E.S. helped to generate astrocyte cultures. B.L.L.C. performed and analyzed all single-cell RNAseq experiments. B.L.L.C. performed the small-molecule screen and all validations. K.C.A. performed all ChIP-seq and ATAC-seq experiments and super-enhancer analysis. B.L.L.C. performed ChIP-seq, ATAC-seq, and bulk RNA-seq data analysis. B.L.L.C. and A.M.S performed and analyzed *in vivo* LPS studies. M.K., E.G., and R.H.M. performed *in vivo* LPC studies and generated electron microscopy images. Y.M.H. analyzed electron microscopy images. B.L.L.C. assembled all figures. B.L.L.C. and P.J.T. wrote the manuscript with input from all authors.

## Competing interests

B.L.L.C. and P.J.T. are listed as inventors on pending patent claims filed by Case Western Reserve University covering methods and compositions for treating neurodegenerative disorders. All other authors declare no competing interests.

---

**Extended Data Figure 1.**
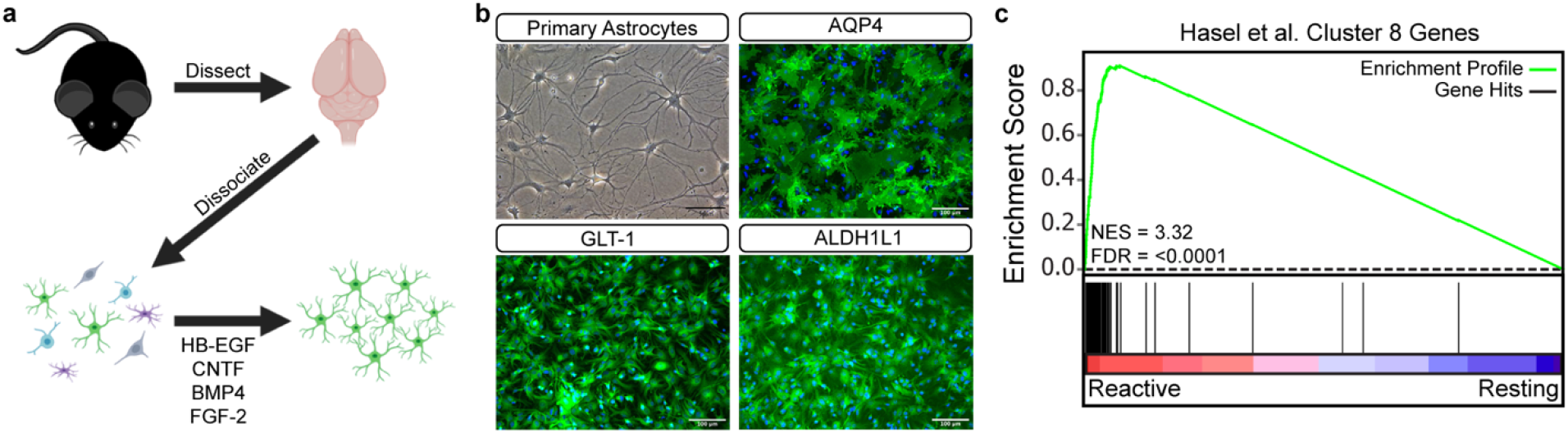
Generation of primary mouse astrocytes and correlation with in vivo reactive astrocyte subtypes. **a**, Overview diagram of astrocyte isolation and enrichment protocol. **b**, Phase contrast image showing prototypical astrocyte morphology. Scale bar is 50um. Immunofluorescence images showing expression of canonical astrocyte markers AQP4, GLT-1 (SLC1A2), and ALDH1L1. Scale bar is 100um. **c**, GSEA comparing pathological reactive astrocytes to Hasel et al. (Nature Neuroscience 2021) Cluster 8 defining genes. Cluster 8 represents a reactive astrocyte state specific to neuroinflammation that also reflects reactive astrocytes in disease models.

**Extended Data Figure 2.**
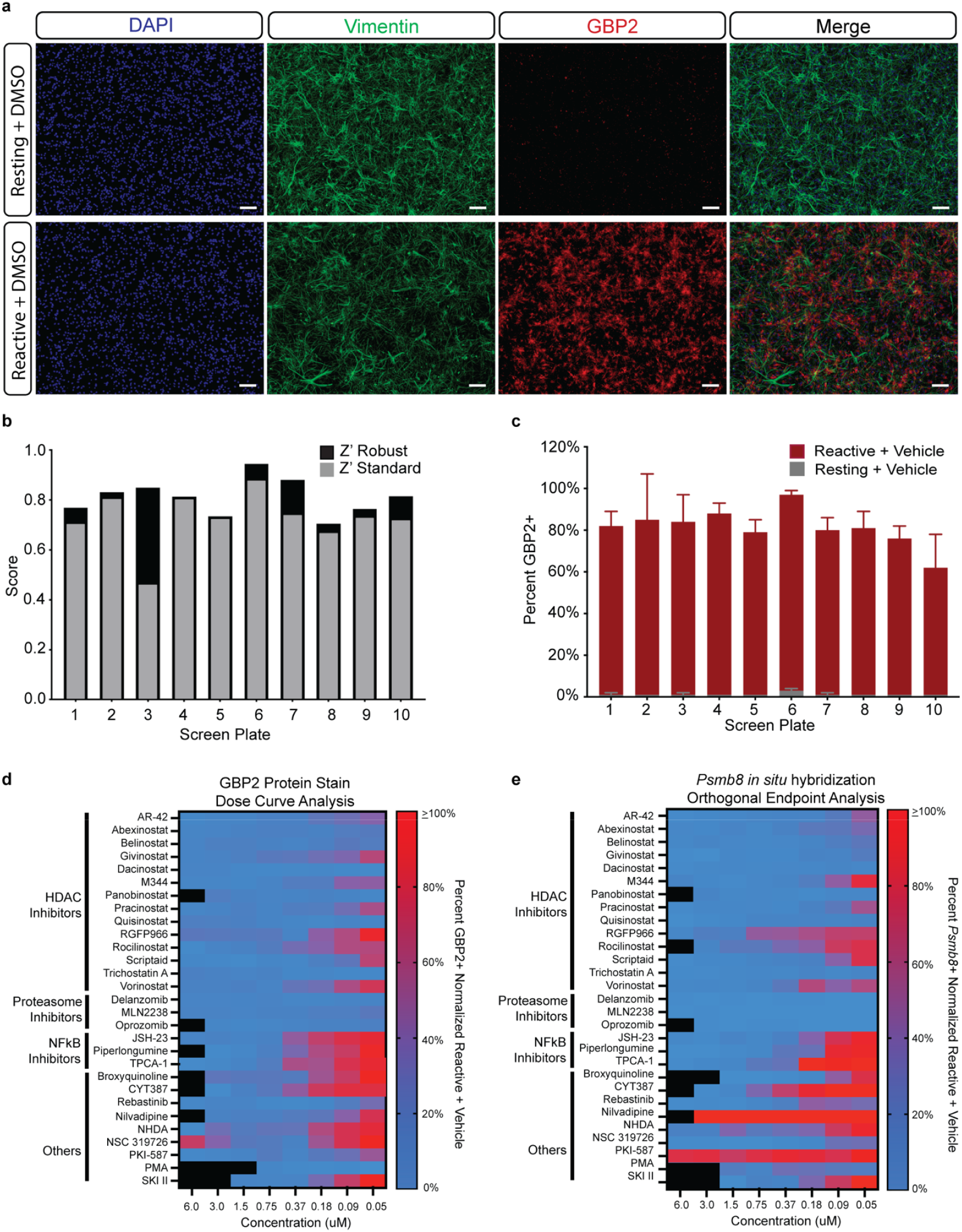
Primary screen quality control and secondary validation of primary hits. **a**, Example images of DMSO vehicle-treated reactive and resting control wells. Scale bar is 100um. **b**, Z-prime standard, and robust scores for each primary screen plate. **c**, Average percent GBP2 positive astrocytes in DMSO vehicle treated reactive and resting control wells on each primary screen plate. **d**, Dose curve analysis of hits from primary screen. Compounds were tested across an 8-point dose curve with decreasing half-steps from 6uM to 0.05uM. Data are presented as the percent of GBP2 positive cells normalized to DMSO vehicle treated reactive astrocyte control wells. *n* = 2 biological replicates (independent astrocyte isolations). Black data points represent toxic doses where total cell number in the well decreased by >50% compared to DMSO vehicle treated reactive astrocyte control wells. **e**, Dose curve analysis of hits from primary screen with *Psmb8* positivity by *in situ* hybridization as a secondary endpoint. Compounds were tested across an 8-point dose curve with decreasing half-steps from 6uM to 0.05uM. Data are presented as the percent of *Psmb8* positive normalized to DMSO vehicle treated reactive astrocyte control wells with an *n* = 1 biological replicate (independent astrocyte isolation). Black data points represent toxic doses where total cell number in the well decreased by >50% compared to DMSO vehicle treated reactive astrocyte control wells.

**Extended Data Figure 3.**
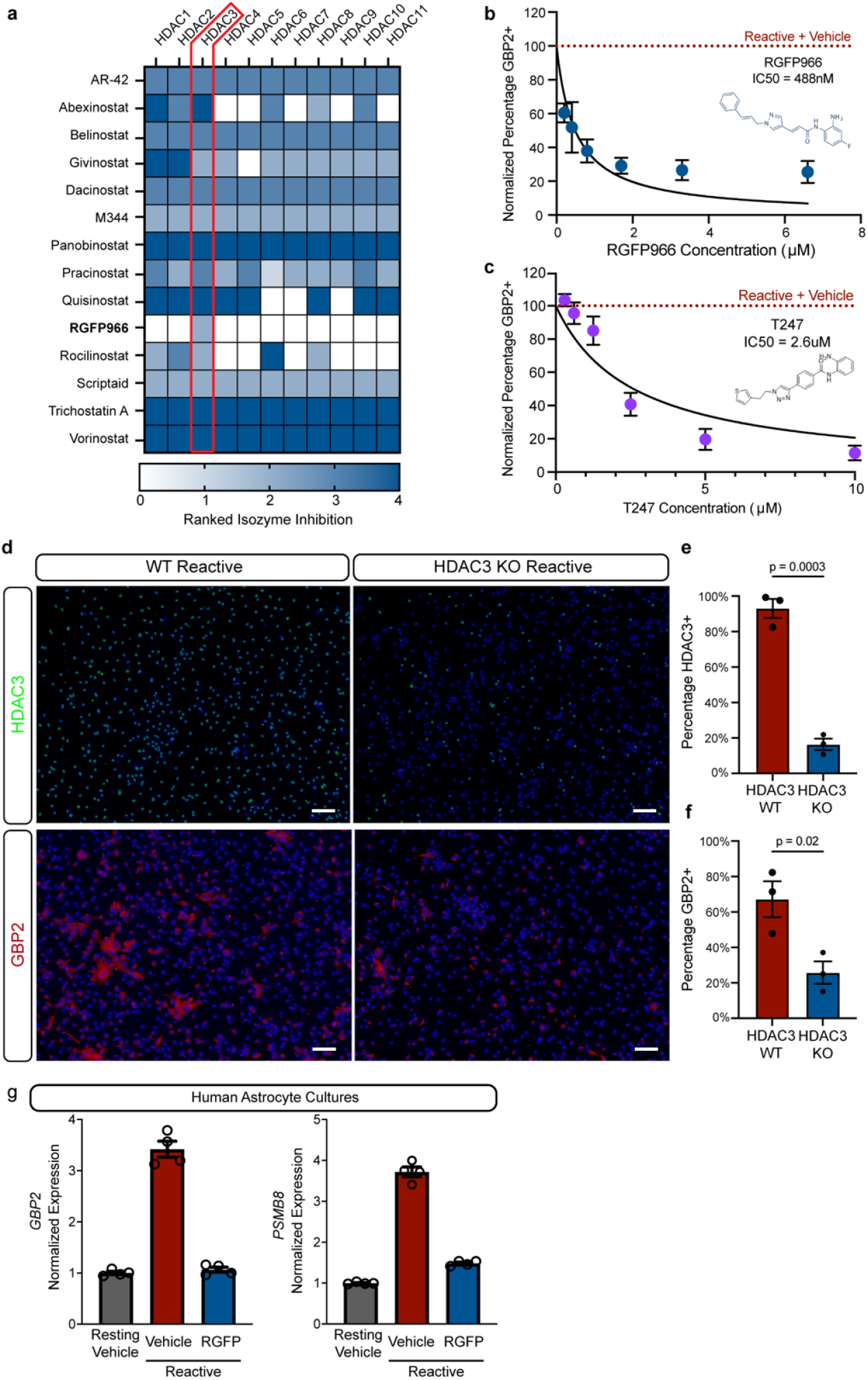
Pharmacological and genetic inhibition of HDAC3 modulates pathological reactive mouse and human astrocytes. **a**, Ranked inhibition against each HDAC isozyme for the validated HDAC inhibitors identified in the primary screen. Highlighted is HDAC3, which is the only shared target between all HDAC inhibitor hits from the primary screen. Ranked efficiency was pulled from target data provided by Selleck Chemical for each compound. **b**, Dose curve and IC50 value for the HDAC3 specific inhibitor RGFP966. **c**, Dose curve and IC50 value for the HDAC3 specific inhibitor T247. **d**, Representative images of wild-type (WT) and HDAC3 knockout (KO) astrocyte cultures exposed to the reactive factors TNF, IL1a, and C1q. Scale bar is 100um. **e**, Quantification of the percentage of cells positive for HDAC3 normalized to WT for the experiment represented in d. Data are presented as the mean ± s.e.m. for *n =* 3 biological replicates (independent astrocyte isolations). p-value generated with a Student’s two-tailed t-test. **f**, Quantification of the percentage of cells positive for GBP2 normalized to WT for the experiment represented in d. Data are presented as the mean ± s.e.m. for *n =* 3 biological replicates (independent astrocyte isolations). p-value generated with a Student’s two-tailed t-test. **g**, *GBP2* and *PSMB8* qPCR results for human iPSC derived resting or reactive (TNF, IL1α, and C1q treated) astrocyte cultures treated with vehicle or 5uM RGFP966. Data are presented as mean ± s.e.m. for *n =* 4 technical replicates.

**Extended Data Figure 4.**
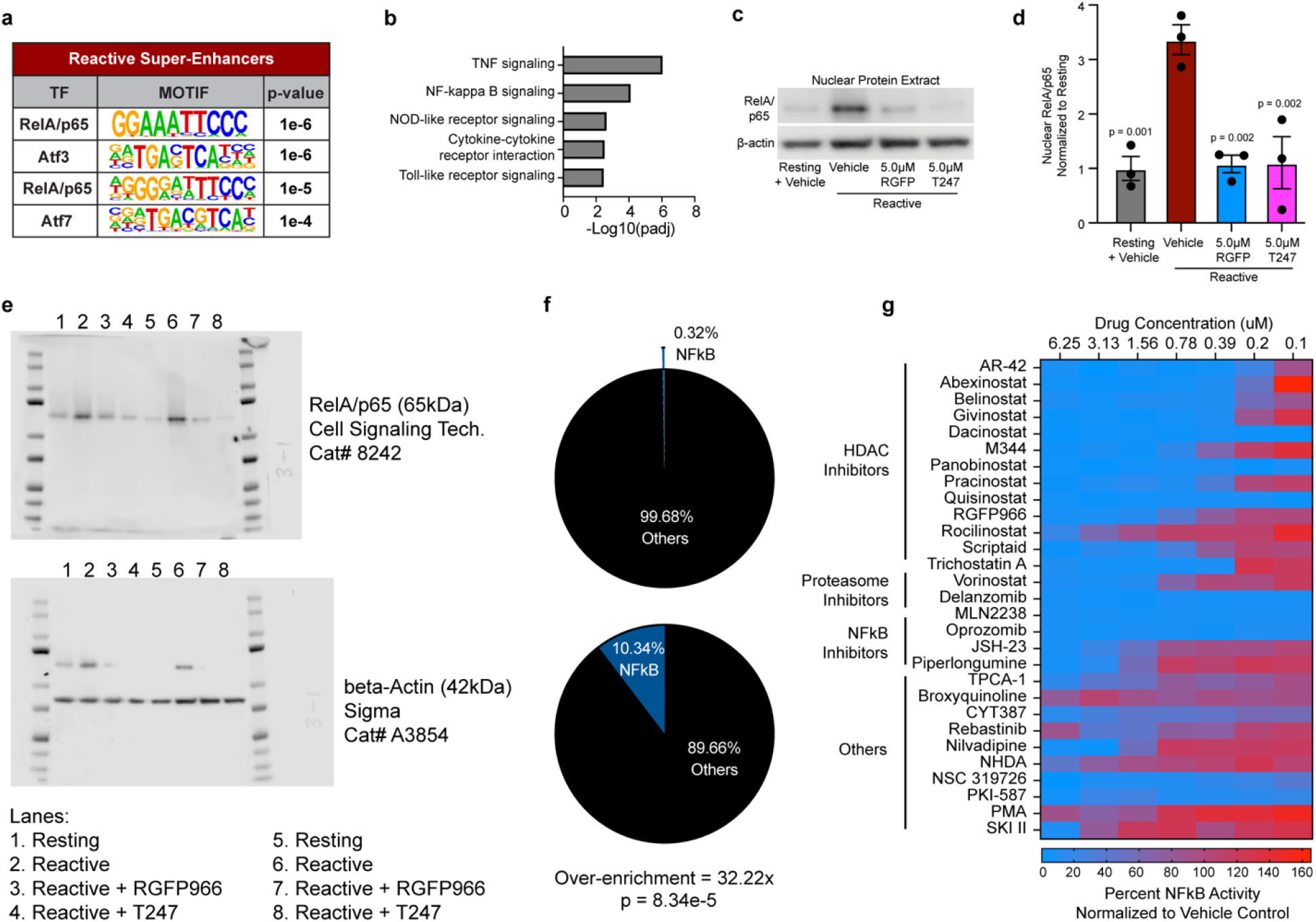
RelA/p65 regulates pathological reactive astrocyte formation and is a unifying target of small-molecule inhibitors of pathological reactive astrocyte formation. **a**, Top results of HOMER transcription factor motif mining beneath pathological reactive astrocyte super-enhancers. **b**, Gene ontology analysis showing KEGG pathways enriched in genes associated with gained pathological reactive astrocyte super-enhancers. **c**, Representative western blot image of nuclear protein extracts probed for RelA/p65 and β-Actin. **d**, Quantification of experiments represented in m. Data are presented as mean ± s.e.m for an *n* = 3 biological replicates (independent astrocyte isolations). p-values generated by a one-way ANOVA with Dunnett post-test for multiple comparisons to reactive plus vehicle control. **e**, Full uncropped western blots that correspond to Fig. 2m. **f**, Distribution of NFkB inhibitors in the full primary screen drug library versus validated hits, showing that NFkB inhibitors are significantly enriched in the validated hit list. p-value generated by a hypergeometric test. **g**, Heatmap showing the normalized NFkB luciferase activity in Jurkat reporter cells treated with the validated hits from the primary drug screen. Data presented as percentage of NFkB activity from a single biological replicate (astrocyte isolation) across an 8-point dose curve from 6.25uM to 0.1uM.

**Extended Data Figure 5.**
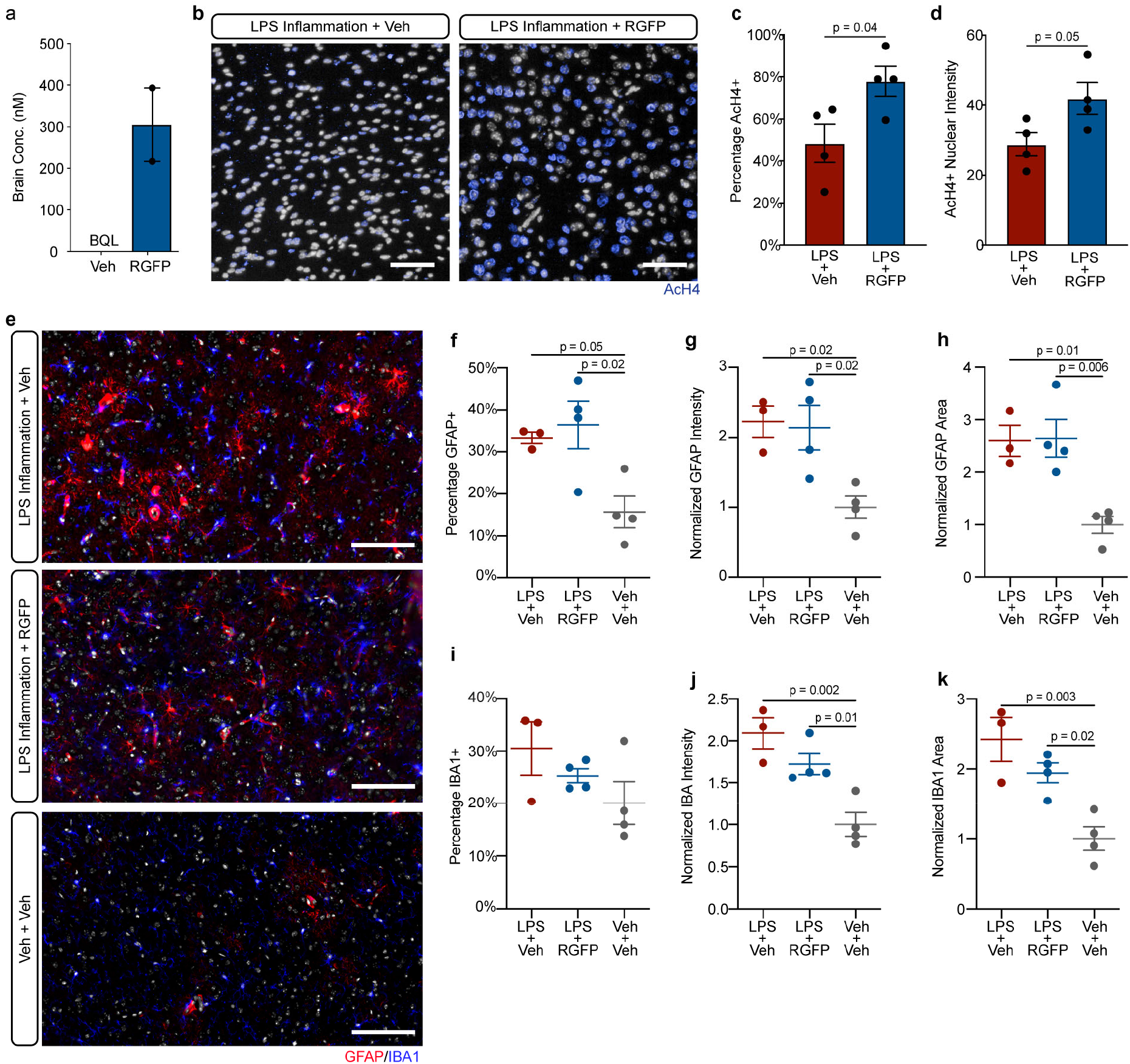
*In vivo* pharmacology of RGFP in mice challenged with systemic LPS to induce neuroinflammation. **a**, Brain concentration of RGFP966 (RGFP) following treatment with vehicle (Veh) or 10mg/kg RGFP. Data presented as mean ± s.e.m. for *n* = 1-2 biological replicates (mice). Concentration of RGFP in brain from vehicle treated mouse was below quantifiable levels (BQL). **b**, Representative images of immunohistochemistry for AcH4 (blue) in the cortex of mice treated chronically with vehicle or 10mg/kg RGFP and then exposed to systemic LPS injections to induce neuroinflammation. Scale bar 100um. **c**, Quantification of the percentage of cells that are AcH4 positive from the experiment represented in b. Data are presented as the mean ± s.e.m. for *n* = 4 biological replicates (mice) per group. p-value generated with a Student’s two-tailed t-test. **d**, Quantification of AcH4 nuclear intensity from the experiment represented in b. Data are presented as the mean ± s.e.m. for *n* = 4 biological replicates (mice) per group. p-value generated with a Student’s two-tailed t-test. **e**, Representative images of immunohistochemistry for GFAP (red) and IBA-1 (blue) in the cortex of mice treated with vehicle or 10mg/kg RGFP and exposed to systemic LPS or saline vehicle. Scale bar 100um. **f-h**, Quantification of GFAP positive astrocytes in the experiment represented in e. Data are presented for the **f**, percentage of GFAP positive cells, **g**, control-normalized GFAP intensity, and **h**, control normalized GFAP area. Data are presented as mean ± s.e.m. for *n* = 3-4 biological replicates (mice) per group. p-value generated by one-way ANOVA with Tukey’s post-test. **i-k**, Quantification of IBA-1 positive microglia in the experiment represented in e. Data are presented for the **i**, percentage of cells IBA-1 positive, **j**, control normalized IBA-1 intensity, and **k**, control normalized IBA-1 area. Data are presented as mean ± s.e.m. for *n* = 3-4 biological replicates (mice) per group. p-value generated by one-way ANOVA with Tukey’s post-test.

**Extended Data Figure 6.**
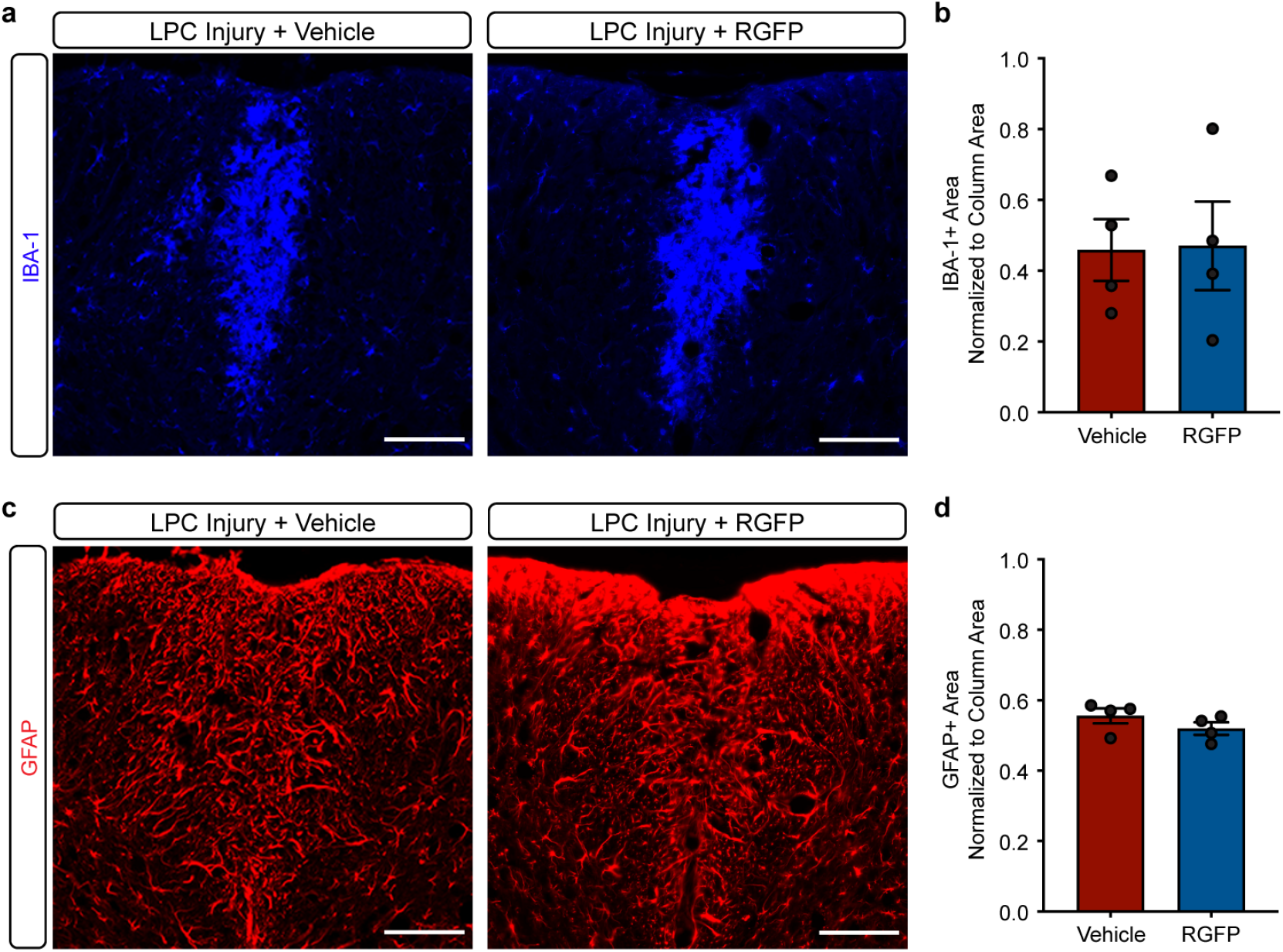
RGFP966 has no effect on generalized gliosis in the toxin-based injury model LPC. **a**, Representative images of LPC lesions from vehicle or RGFP966 (RGFP) treated mice stained for IBA-1. Scale bar is 100um. **b**, Quantification of the IBA-1 positive area divided by the dorsal column area from staining represented in a. Data are presented as mean ± s.e.m. for *n* = 4 biological replicates (mice) for each group. **c**, Representative images of LPC lesion from vehicle or RGFP treated mice stained for GFAP. Scale bar is 100um. **d**, Quantification of the GFAP-positive area divided by the dorsal column area from staining represented in a. Data are presented as mean ± s.e.m. for *n* = 4 biological replicates (mice) per group.

